## Supplementary figures and images for "Chemotherapy resistance due to epithelial-to-mesenchymal transition is caused by abnormal lipid metabolic balance"

### Figure 1 figure supplement 1

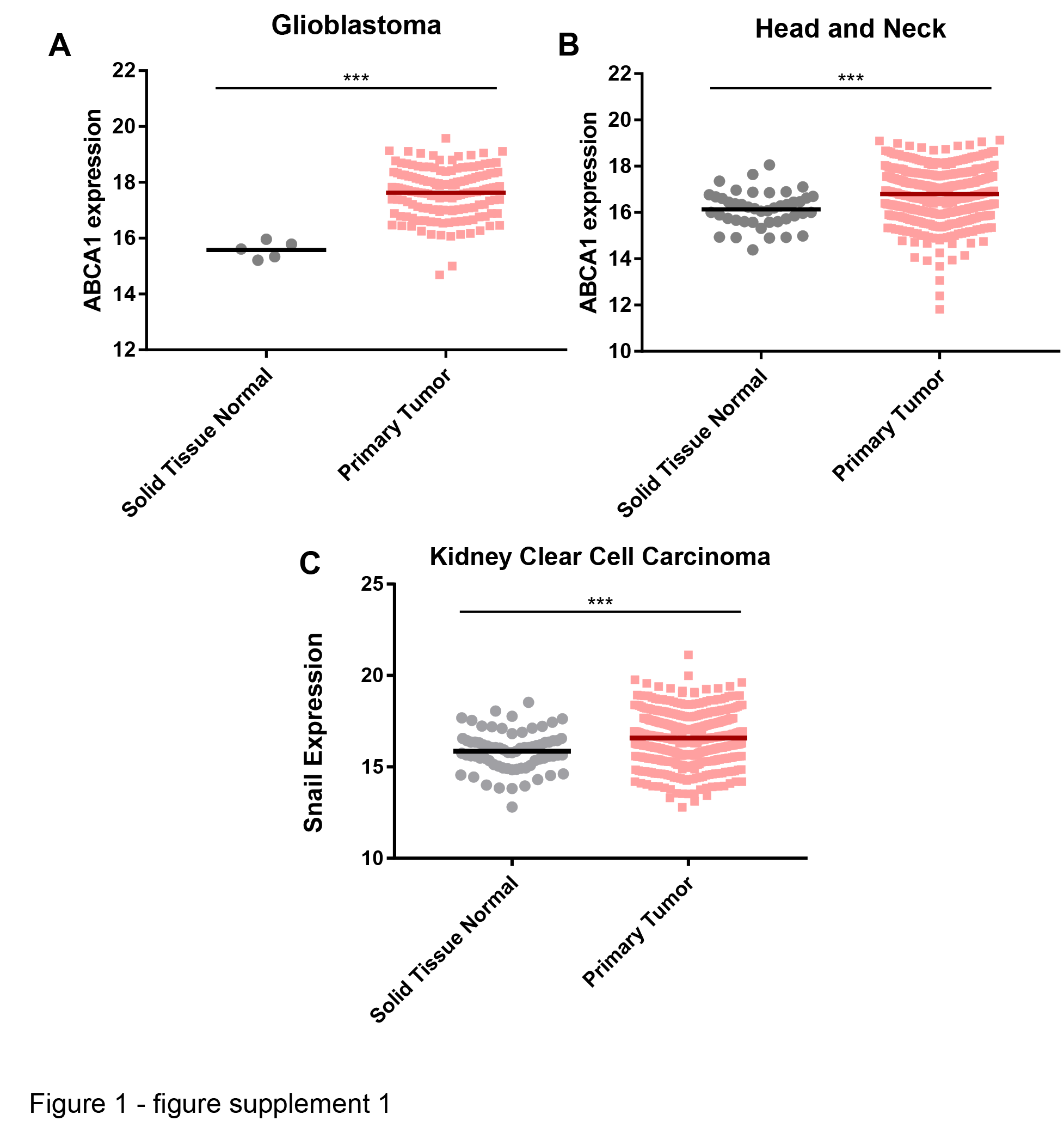

### Figure 1 figure supplement 2

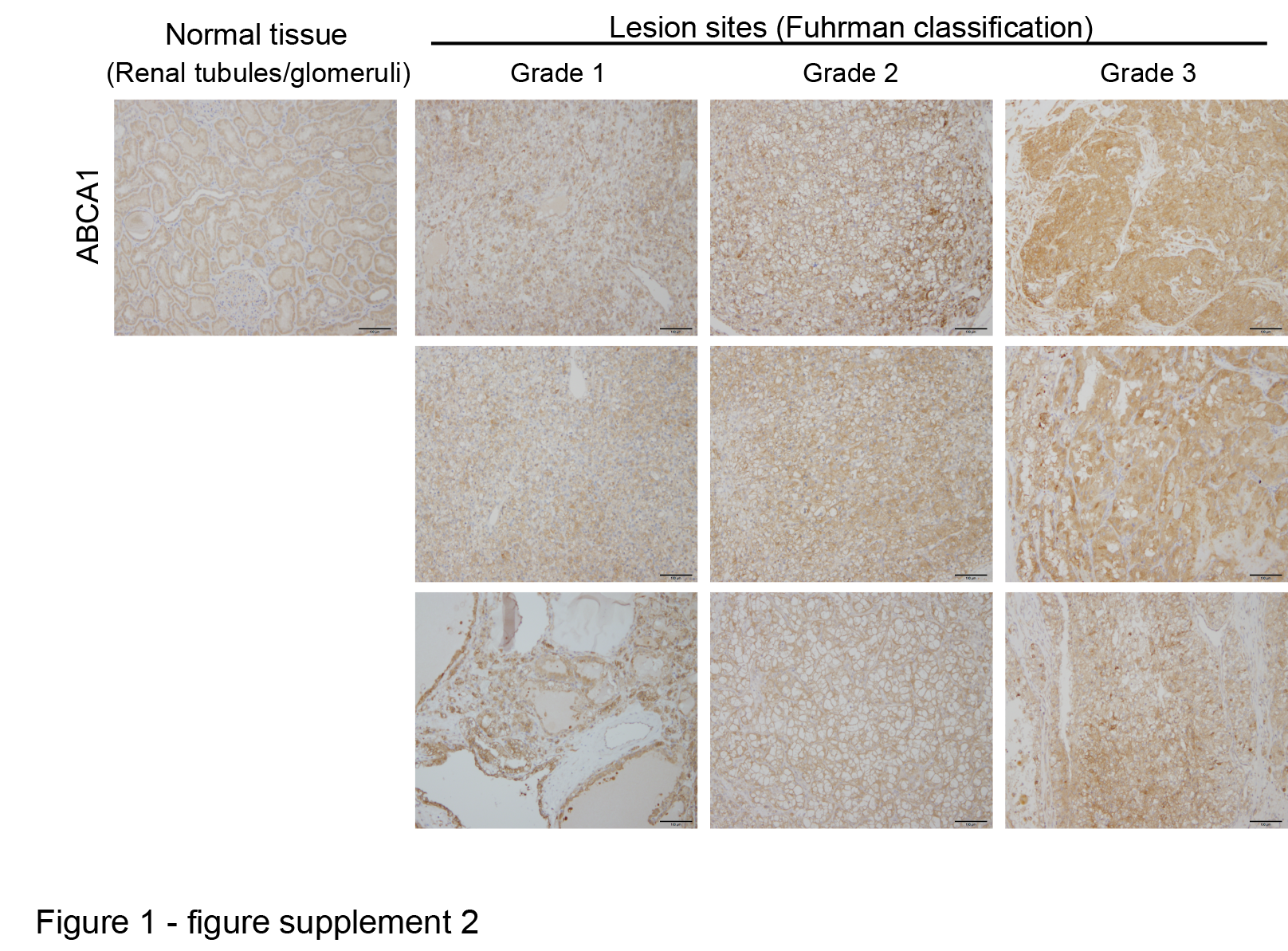

### Figure 2 figure supplement 1

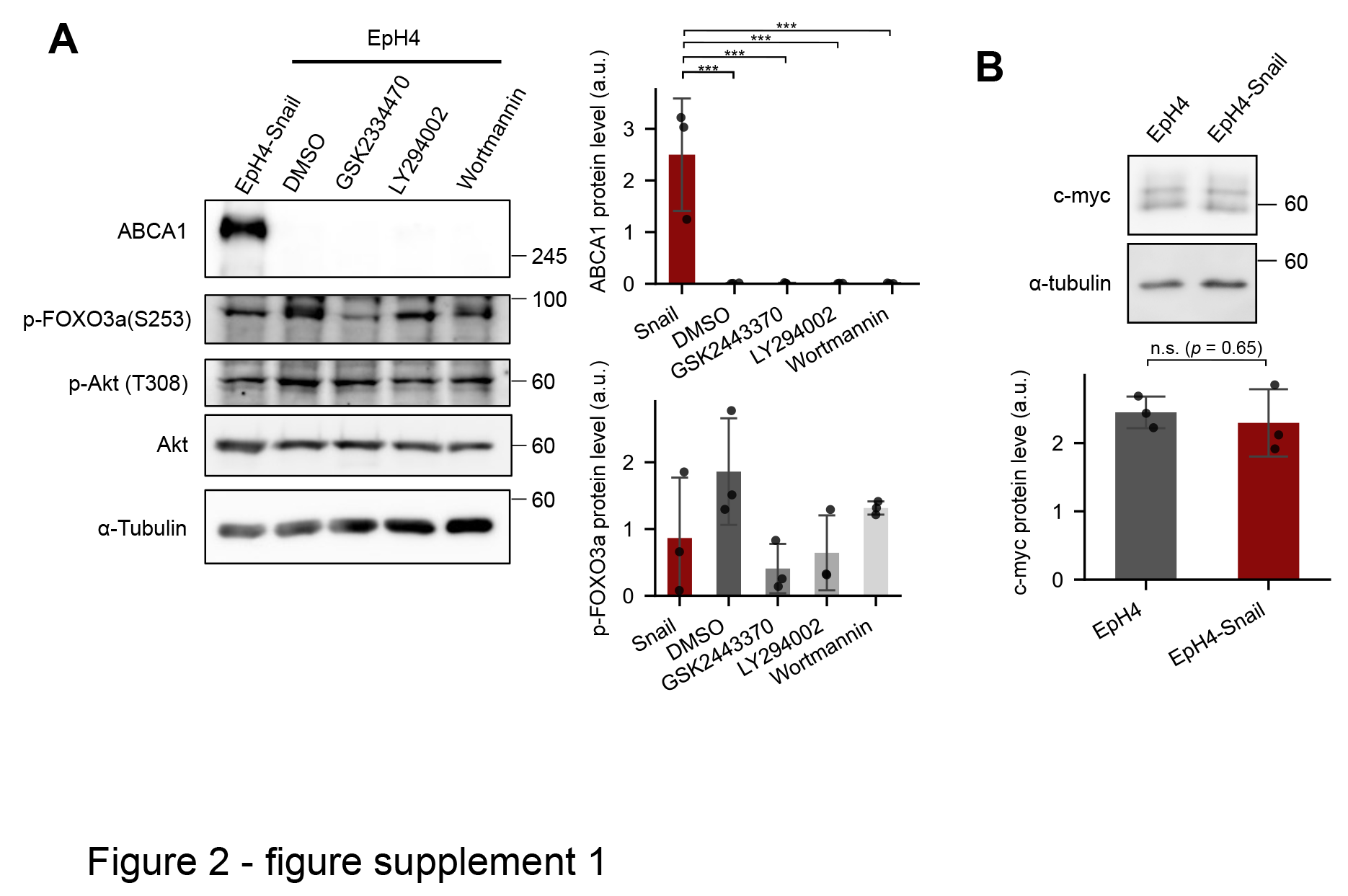

### Figure 2 figure supplement 2

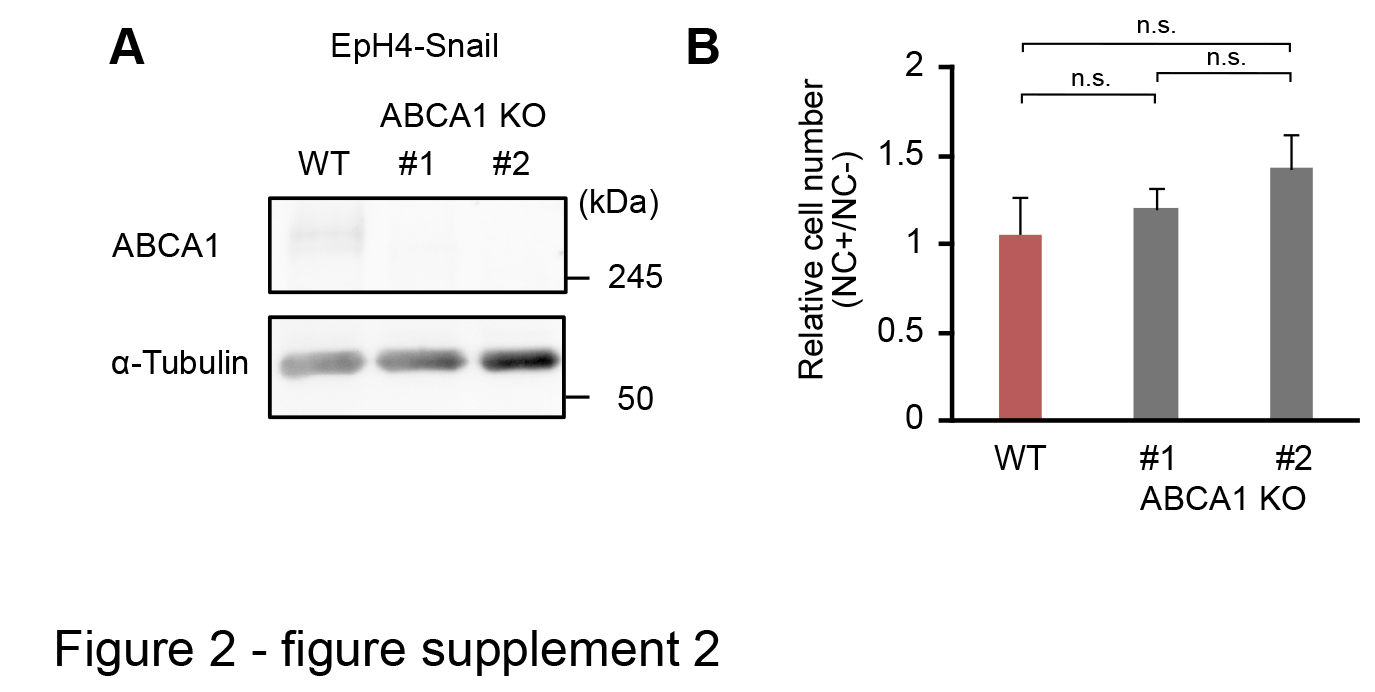

### Figure 2 figure supplement 3

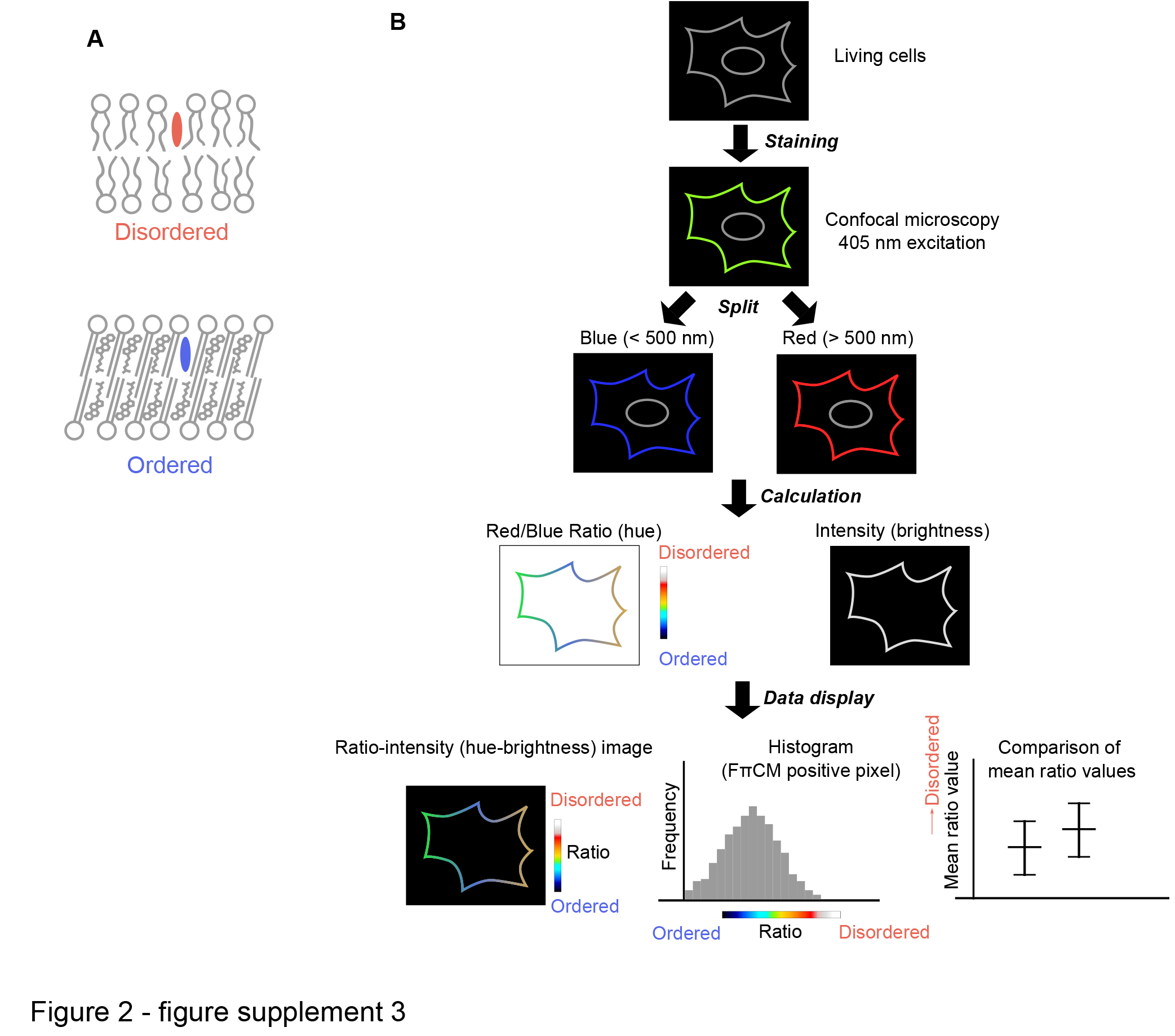

### Figure 3 figure supplement 1

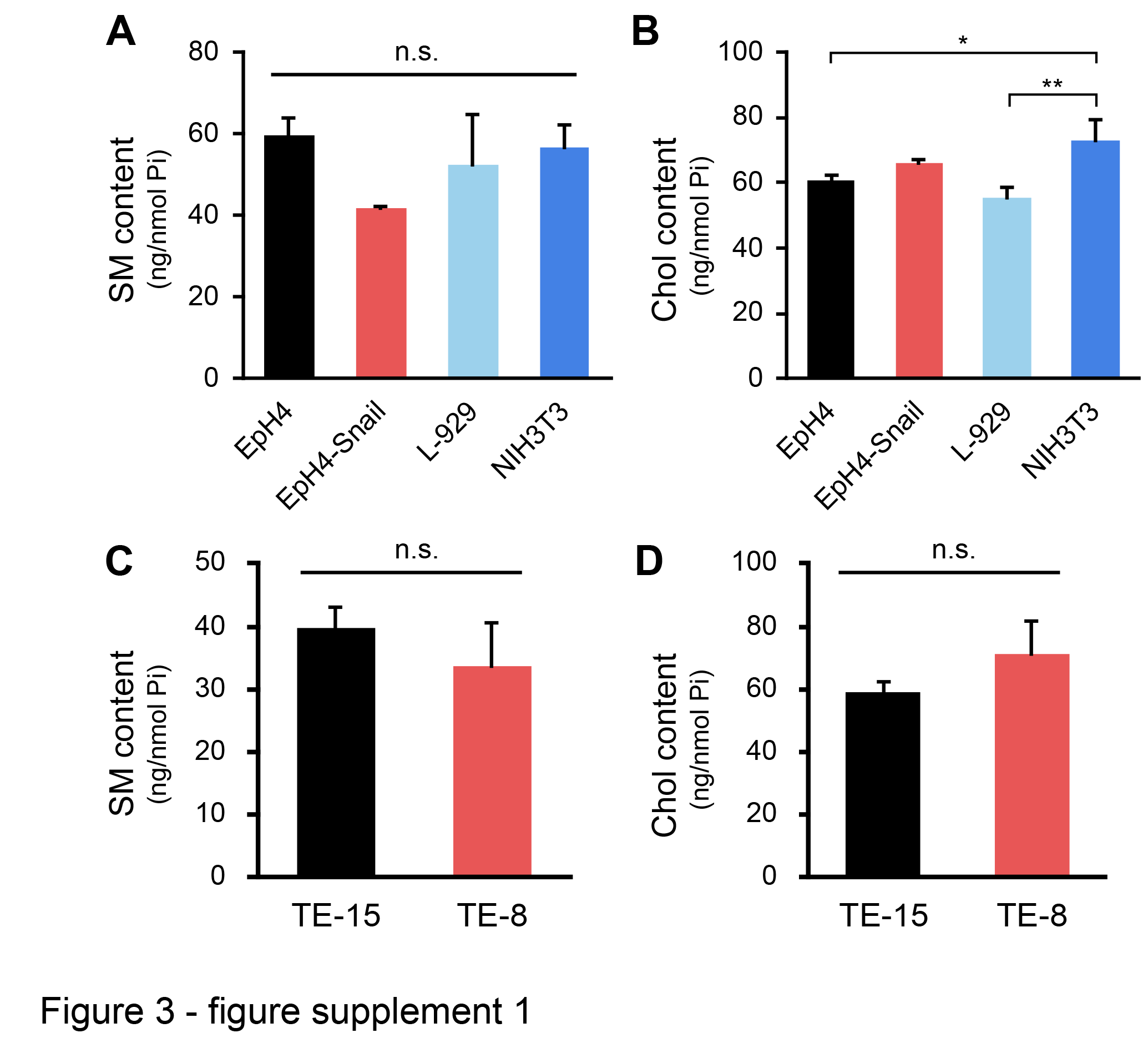

### Figure 4 figure supplement 1

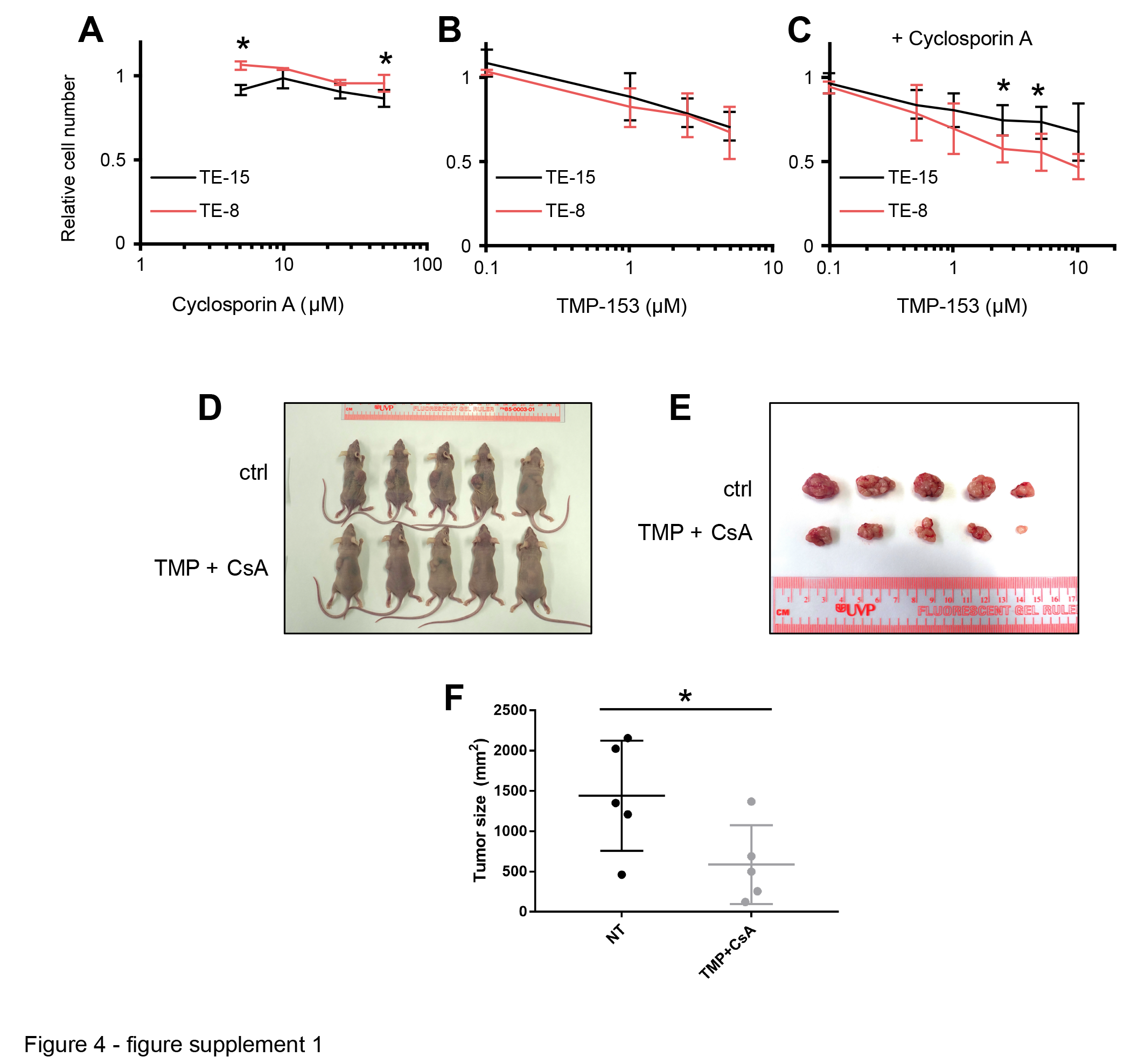
