## Supplementary material for "Chemotherapy resistance due to epithelial-to-mesenchymal transition is caused by abnormal lipid metabolic balance": Table S1

**Table S1 Primer sequences used in this study**

| **Gene** | | **Primer sequence** | |
| --- | --- | --- | --- |
| **CerS isozymes** | |  |  |
| **mouse** | *CerS1* | forward | 5'-CTGTACATTGTGGCTTTCGC-3' |
|  |  | reverse | 5'-GGCTGTCTGAGCTTCCAGAG-3' |
| **mouse** | *CerS2* | forward | 5'-CCCTGCTCTTTCTCGTCATTC-3' |
|  |  | reverse | 5'-CGTGACAAAAGGTCTACCTCC-3' |
| **mouse** | *CerS3* | forward | 5'-TTTCAAGCATTCCACAAGCAAAC-3' |
|  |  | reverse | 5'-ATCGCCAGCAAGACTCCT-3' |
| **mouse** | *CerS4* | forward | 5'-GGCTGTGCGAATTGTCTT-3' |
|  |  | reverse | 5'-CGGGTTGGGCTTTATCTTT-3' |
| **mouse** | *CerS5* | forward | 5'-AGTAAGGTTATTGGCTCTGTGTT-3' |
|  |  | reverse | 5'-GCTCAGAAGTGACCAGACAAT-3' |
| **mouse** | *CerS6* | forward | 5'-CCGGAGTGGACAAAGCAAGAT-3' |
|  |  | reverse | 5'-GCCGTTCTTACCTCTCTGGAAGATG-3' |
| **Elovl isozymes** | |  |  |
| **mouse** | *Elovl1 variant1* | forward | 5'-TGGAGGCTGTTGTGAACTTG-3' |
|  |  | reverse | 5'-ATGAAGCCTCGAAGTTGGAA-3' |
| **mouse** | *Elovl1 variant2* | forward | 5'-GATCCCTTTG AACCCTTCAC-3' |
|  |  | reverse | 5'-TCAGAAACCGCATGCTTCAT-3' |
| **mouse** | *Elovl2* | forward | 5'-GATCCCTTTGAACCCTTCAC-3' |
|  |  | reverse | 5'-GAGAATCTCGTGGTCCAAACA-3' |
| **mouse** | *Elovl3* | forward | 5'-CGGGTTAAAAATGGACCTGA-3' |
|  |  | reverse | 5'-CCAACAACGATGAGCAACAG-3' |
| **mouse** | *Elovl4* | forward | 5'-ACCGTGGAGTTCTATCGCTG-3' |
|  |  | reverse | 5'-TGCTTATGCTTATCGTTGGC-3' |
| **mouse** | *Elovl5* | forward | 5'-CTTGCACATCCTCCTGCTC-3' |
|  |  | reverse | 5'-CTGAGTGACGCATCGAAATG-3' |
| **mouse** | *Elovl6* | forward | 5'-CCAATGGATGCAGGAAAACT-3' |
|  |  | reverse | 5'-AACTTGGCTCGCTTGTTCAT-3' |
| **mouse** | *Elovl7* | forward | 5'-GGAAAATGGCGTTCAGTGAT-3' |
|  |  | reverse | 5'-CATAGAGGCCCAGGATGATG-3' |
| **quantitative analysis** | |  |  |
| **mouse** | *ZO-1* | forward | 5’-AGGTGGGTCGGAATTCGGAGATAGTGTGGGTTTGCGAC-3’ |
|  |  | reverse | 5’-GATTGTCGACGAATTCTCATAACTGTTCAGCTCTGTTCTTAT-3’ |
| **mouse** | *Elovl7* | forward | 5'-ATGGCGTTCAGTGATCTTACATCG-3' |
|  |  | reverse | 5'-CCTGCAGCAAATTTGACTCCAAAC-3' |
| **mouse** | *CerS3* | forward | 5'-CGTTTAGAAAATGGTTCTGGTCGG-3' |
|  |  | reverse | 5'-GTTACACTTCTTTGCCAGTCC-3' |
